# DNA origami as a nanomedicine for targeted rheumatoid arthritis therapy through reactive oxygen species and nitric oxide scavenging

**DOI:** 10.1101/2021.12.30.474611

**Authors:** Yuxuan Ma, Zhangwei Lu, Ye Shi, Zhe Li

## Abstract

High levels of reactive oxygen species (ROS) and nitric oxide (NO) generated by M1 macrophages induce inflammation in the development of rheumatoid arthritis (RA). The eliminating of ROS and NO therefore represents an alternative strategy for RA treatment. Because DNA molecules possess ROS- and endogenous NO-scavenging capability, herein, we develop a nanomedicine based on triangular DNA origami nanostructures for targeted RA treatment. We showed that folic acid-modified triangular DNA origami nanostructures (FA-tDONs) could reduce ROS and NO simultaneously inside proinflammatory M1 macrophages, leading to their polarization into anti-inflammatory M2 subtype. Further in vivo studies confirmed that FA-tDONs could actively target inflamed joints in collagen-induced arthritis (CIA) mice, attenuate inflammatory cytokines and alleviate disease progression. This work demonstrated that DNA origami itself could act as a potential nanomedicine for targeted RA treatment.

## Introduction

Rheumatoid arthritis (RA) is a chronic inflammatory autoimmune disorder that affects ∼1% of the adult population by persistent synovitis and irreversible joint disability.^1, 2^ Currently, synthetic disease-modifying antirheumatic drugs (DMARDs), e.g., methotrexate, are the optimal approach for early RA treatment. ^3^However, due to unknown action models and serious side effects, some patients respond insufficiently, and long-term disease remission is not achieved.^4^ The combination of biological DMARDs such as TNF-α inhibitors with methotrexate might be a better choice for low disease activity. However, patient response rates are only 30%∼40%, accompanied by adverse reactions such as infections and tuberculosis.^5, 6^ Hence, developing novel therapeutic strategies for RA is in urgent demand.

One of the hallmarks of RA is persistent synovitis that results from the sustained influx of immune cells into the joints.^7^ In particular, proinflammatory M1 macrophages are predominant in RA joints and play a crucial role in the promotion of monocyte recruitment, proliferation of synovial fibroblasts and formation of osteoclasts by secreting inflammatory mediators such as IL-1β and TNF-α.^8, 9^ In contrast, anti-inflammatory M2 macrophages secrete anti-inflammatory cytokines, such as IL-10.^8, 10^ The balance of M1 and M2 macrophages is affected by the levels of reactive oxygen species (ROS) and nitric oxide (NO).^11, 12^ While high levels of ROS and NO induce polarization of macrophages to the M1 subtype, the scavenging of ROS/NO can switch M1 macrophages into M2 macrophages and therefore represents an alternative strategy for RA treatment.^13^ Though a number of nanodrugs (e.g. manganese ferrite/ceria nanoparticle,^14^ silver nanoparticle^15^ and NO-scavenging hydrogel^16, 17^) that can either eliminate ROS or NO have been developed to treat RA, there is still deficiency of nanomedicines with fully predictable structures, minima off-target toxicity, long duration and stability.

With high programmability and biostability, DNA nanostructures have been widely used in drug delivery and biosensing.^18, 19^ Recent studies have also shown that DNA nanostructures possess good ROS- and endogenous NO-scavenging capability.^20-22^ Taking all the advantages, herein, we developed a DNA nanostructure-based nanomedicine for RA treatment. Specifically, we designed a triangular DNA origami nanostructure with folic acid modification (FA-tDONs) to actively target arthritis sites and studied their therapeutic potential in an RA mouse model (Scheme 1). We have demonstrated that the biocompatible FA-tDONs were effective in regulating the M1/M2 switch by simultaneously scavenging ROS and endogenous NO, and alleviating inflammation of the RA joints. This work demonstrated that DNA origami nanostructure itself could act as a potential nanomedicine for RA treatment.

**Scheme 1.**
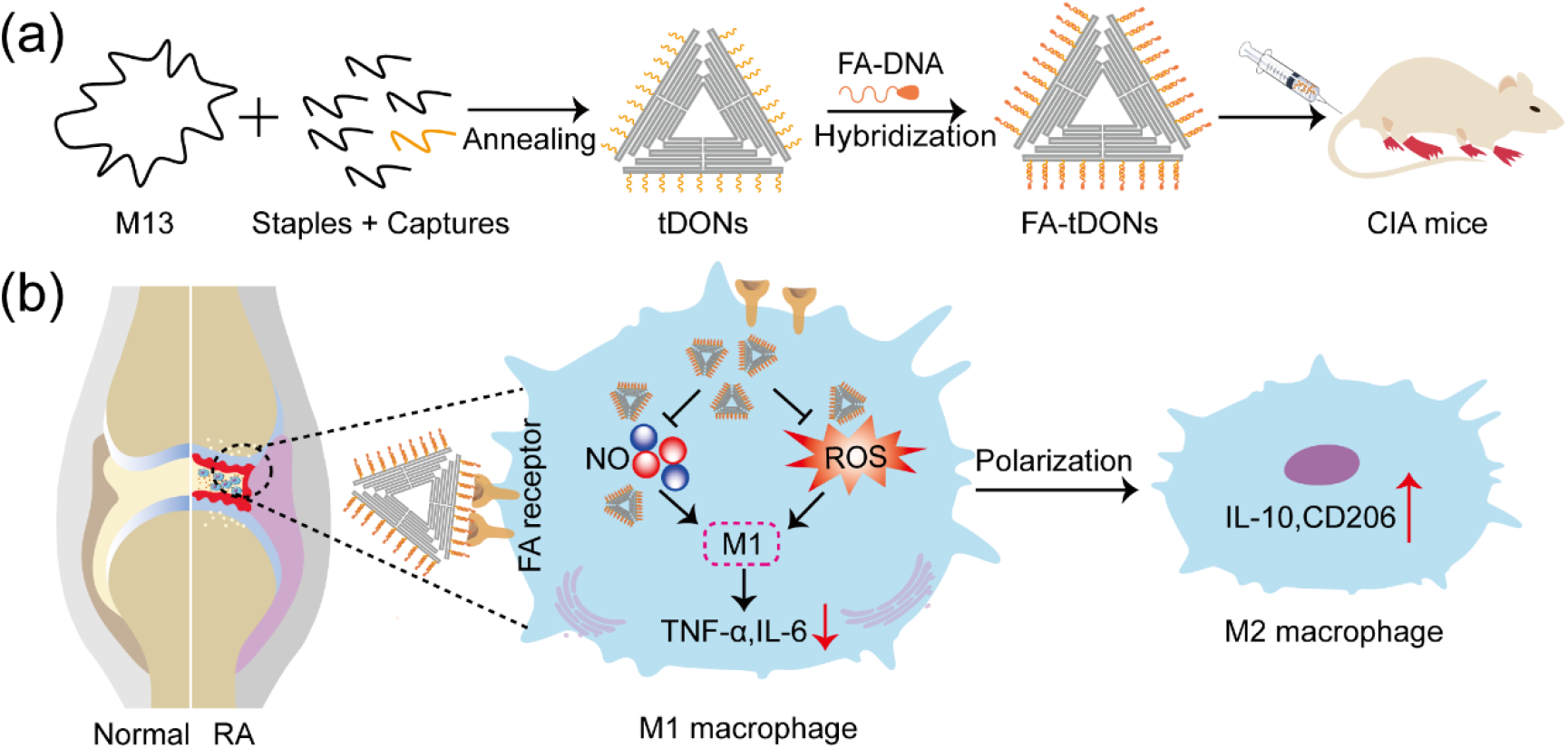
Design of FA-tDONs nanomedicine. (a) Schematic illustration showing the composition of FA-tDONs, in which triangular DNA origami nanostructures (tDONs) are modified with folic acid (FA), a targeting ligand to M1 macrophages. (b) FA-tDONs can actively target M1 macrophages and polarize proinflammatory M1 macrophages into anti-inflammatory M2 macrophages by simultaneously scavenging ROS and NO for RA therapy.

## Results and discussion

### Preparation and Characterization of FA-tDONs

First, a 90-nm triangular DNA origami nanostructure (tDON) was designed, which could be passively accumulated at the RA sites due to the “ELVIS”^23^ (extravasation through leaky vasculature and subsequent inflammatory cell-mediated sequestration) effect. Next, to achieve actively targeted delivery, folic acid (FA), which shows high affinity for folate receptor beta (FR-β) overexpressed on M1 macrophages^24^ in RA, was modified onto the surface of triangular DNA origami nanostructures. The FA was conjugated to an oligo strand (FA-oligo) through an amide reaction between FA-NHS and the amine group of the oligo (figure S1), and the conjugation was confirmed by mass spectrometry and native gel characterizations (figure S1 and S2). FA-oligo was then introduced onto the surface of tDONs via DNA hybridization, leading to the formation of FA-modified tDONs (FA-tDONs). The assembled nanostructures showed clear master bands on agarose gel, and the migration rate of tDONs was slightly faster than that of FA-tDONs (Figure 1a), indicating their successful assembly. The morphology and size of FA-tDONs were further characterized by atomic force microscopy (AFM) and dynamic light scattering (figure 1b, 1c and S3), which showed no obvious difference from tDONs as expected. The stability of tDONs and FA-tDONs against 10% fetal bovine serum (FBS) after 24 h incubation was tested, and both structures remained intact (figure 1d), suggesting their stability in biological fluid.

**Figure 1.**
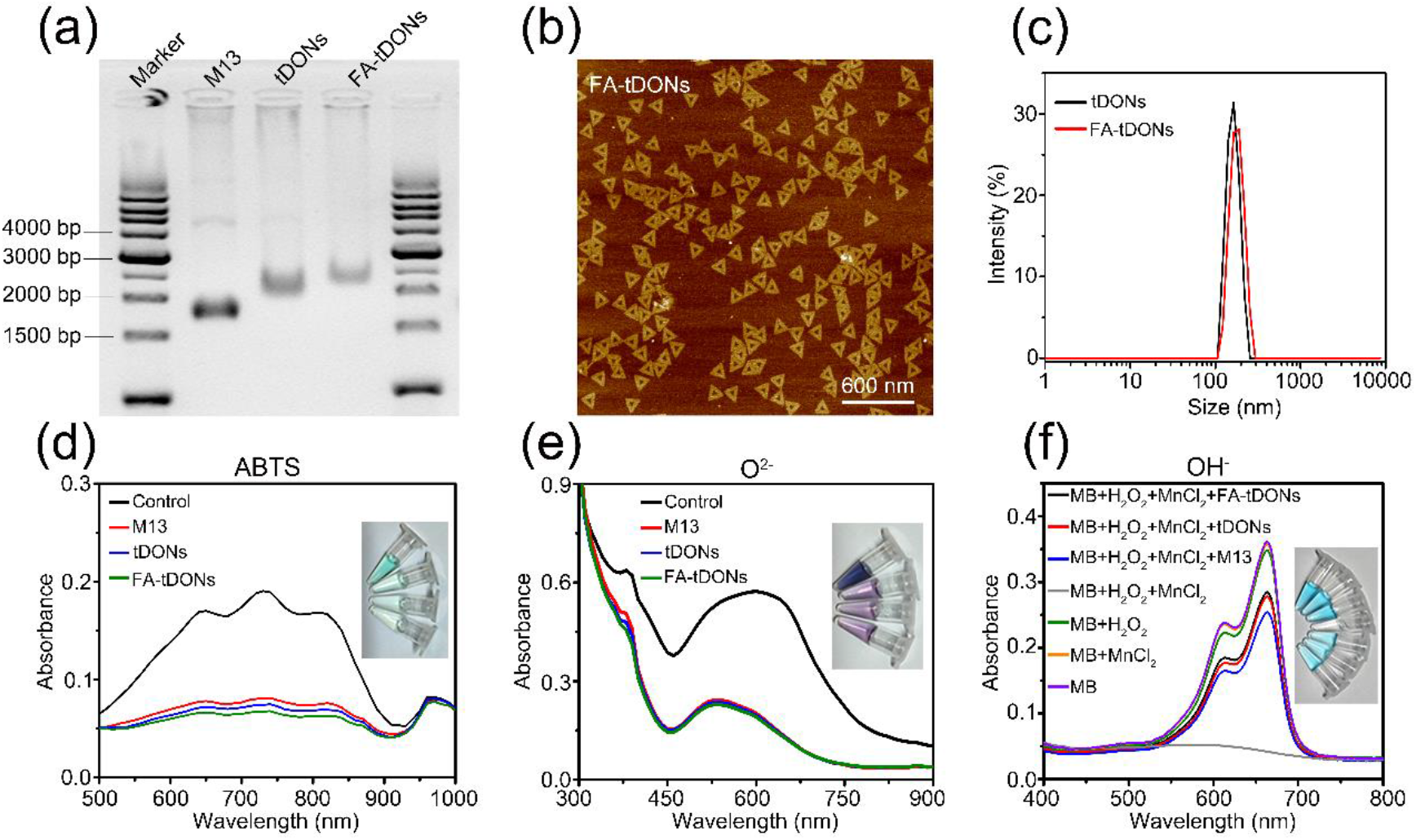
Characterization of FA-tDONs. (a) 1% agarose gel analysis of M13, tDONs and FA-tDONs. (b) Representative AFM images of FA-tDONs. (c) Hydrodynamic sizes of tDONs and FA-tDONs. (d) ABTS radical scavenging activity of M13 ssDNA, tDONs and FA-tDONs for 12 h. The characteristic absorbance of ABTS radicals at 734 nm decreased after incubation with DNA materials. (e) Superoxide radical scavenging activity of M13 ssDNA, tDONs and FA-tDONs in the NADH-PMS-NBT system. (f) Hydroxyl radical scavenging activity of M13 ssDNA, tDONs and FA-tDONs probed by MB.

The ROS scavenging effect of FA-tDONs was evaluated in testing tubes. First, we measured the radical scavenging ratios of M13 single-stranded DNA, tDONs, and FA-tDONs to model stable free radicals (ABTS). Under the same number of DNA bases, all groups showed 50% elimination efficiencies when incubated with ABTS at 12 h (Figure 1d and S5). In addition, the scavenging ability of FA-tDONs for superoxide radicals (O_2_ ^−^•) and hydroxyl radicals (•OH), two typical types of ROS commonly seen in living organisms, was further characterized. The characteristic absorbance of blue chromagen formazan produced by the reaction of O_2_ ^−^• with NBT was monitored, where a lower absorbance indicated a stronger O_2_ ^−^• scavenging ability^25^. The •OH scavenging ability was detected by the characteristic absorbance of MB, which can be rapidly degraded by •OH, accompanied by the disappearance of UV–vis adsorption. FA-tDONs and tDONs exhibited strong scavenging activity toward both O_2_ ^−^• and •OH (Figure 1e and 1f). In summary, these results demonstrate that FA-tDONs have high radical scavenging activities.

### In vitro cellular uptake

We used the macrophage cell line RAW264.7 as a model. According to previous reports, lipopolysaccharide (LPS) stimulation can activate RAW264.7 cells to the M1 phenotype and promote the secretion of proinflammatory cytokines. LPS (10 μg/mL) effectively activated RAW264.7 cells by significantly increasing cytokines such as TNF-α and IL-6 (Figure S4). Confocal images revealed that the activated macrophages exhibited high red fluorescence upon treatment with Cy5 labeled FA-tDONs compared to tDONs, indicating significant promotion of cellular internalization by modification with FA on the surface of DNA origami nanoparticles (figure 2a, upper and middle lines). Additionally, after pretreatment of activated macrophages with free FA to block the FR-β pathway, the fluorescent signal of FA-tDONs was hardly observed, which further confirmed the contribution of FR-β to FA-tDON internalization (figure 2a, bottom line). However, the normal macrophages (without LPS stimulation) showed weak red fluorescence after the incubation with FA-tDONs (figure 2b). Consistently, flow cytometry analysis further demonstrated the targeting ability of FA-tDONs to active macrophages (Figure 2c and 2d).

**Figure 2.**
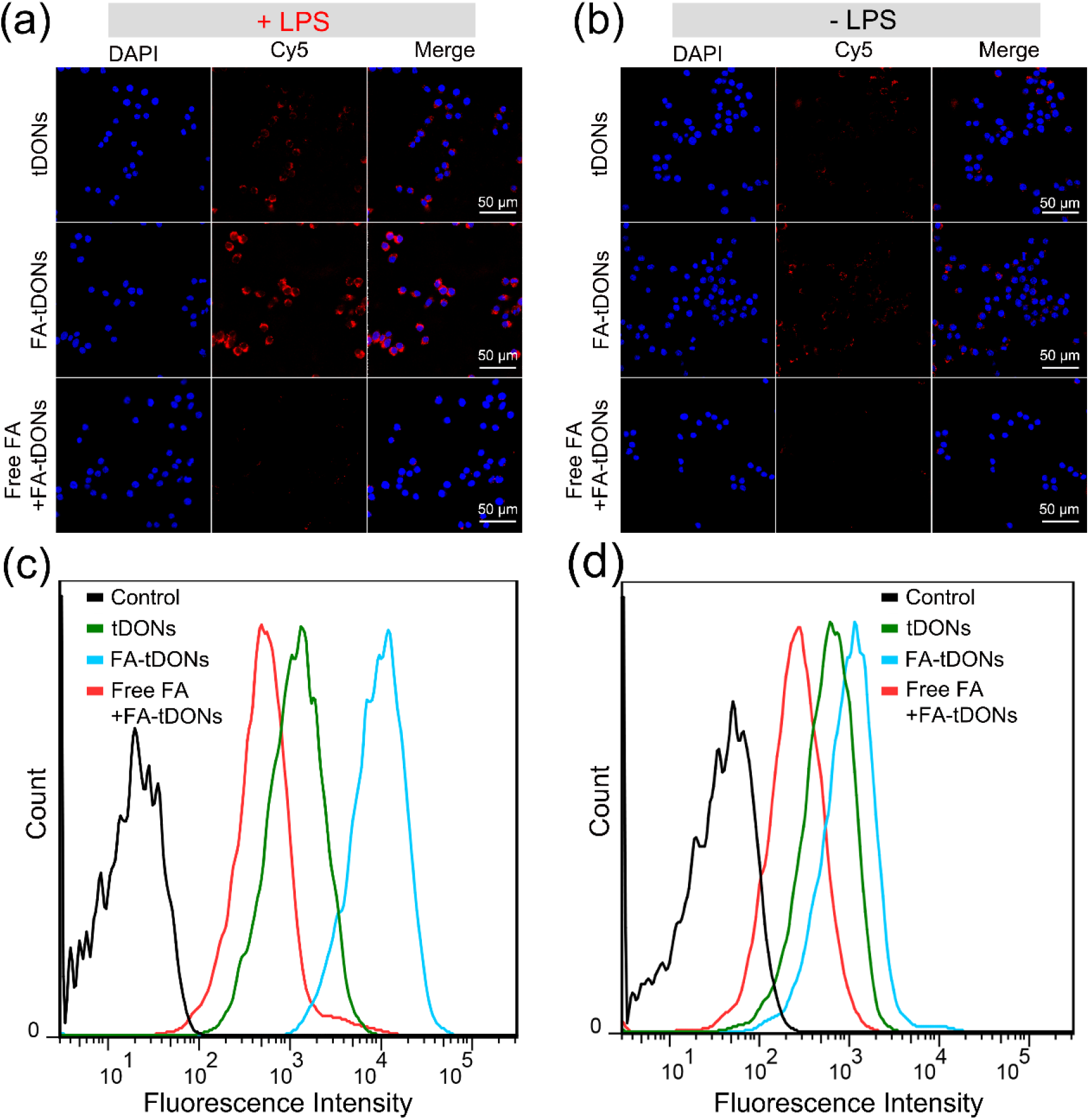
The cellular uptake efficiency of FA-tDONs by M1 macrophages. Confocal laser fluorescence imaging of (a) LPS-activated RAW264.7 cells and (b) RAW264.7 cells without LPS stimulation treated with Cy5-labeled tDONs and FA-tDONs (in the absence or presence of 1 mM free FA) for 2 h. (c)(d) Corresponding flow cytometry analysis.

### In vitro ROS and endogenous NO scavenging ability of FA-tDONs

To visualize the ROS scavenging activity of FA-tDONs in activated macrophages, 2′-7′-dichlorodihydrofluorescein diacetate (DCFH-DA) was used to detect the total intracellular ROS levels. After treatment with LPS, a strong fluorescence signal of DCFH-DA was observed in the activated macrophages compared to the normal RAW264.7 cells without LPS stimulation (figure 3a). The fluorescence intensity of DCFH-DA significantly decreased after the addition of tDONs or FA-tDONs, indicating their efficient removal of ROS. In addition, the relative fluorescence intensity of each group was measured by flow cytometry for quantitative comparison (figure 3b and 3c). Consistently, the fluorescence intensity increased ∼60-fold after LPS stimulation, indicating the activation of macrophages, and decreased ∼40- and 80-fold after treatment with tDONs and FA-tDONs for 24 h respectively, suggesting efficient ROS scavenging activity. Further investigation showed that tDONs and FA-tDONs scavenged ROS in a concentration-dependent manner (figure S6).

**Figure 3.**
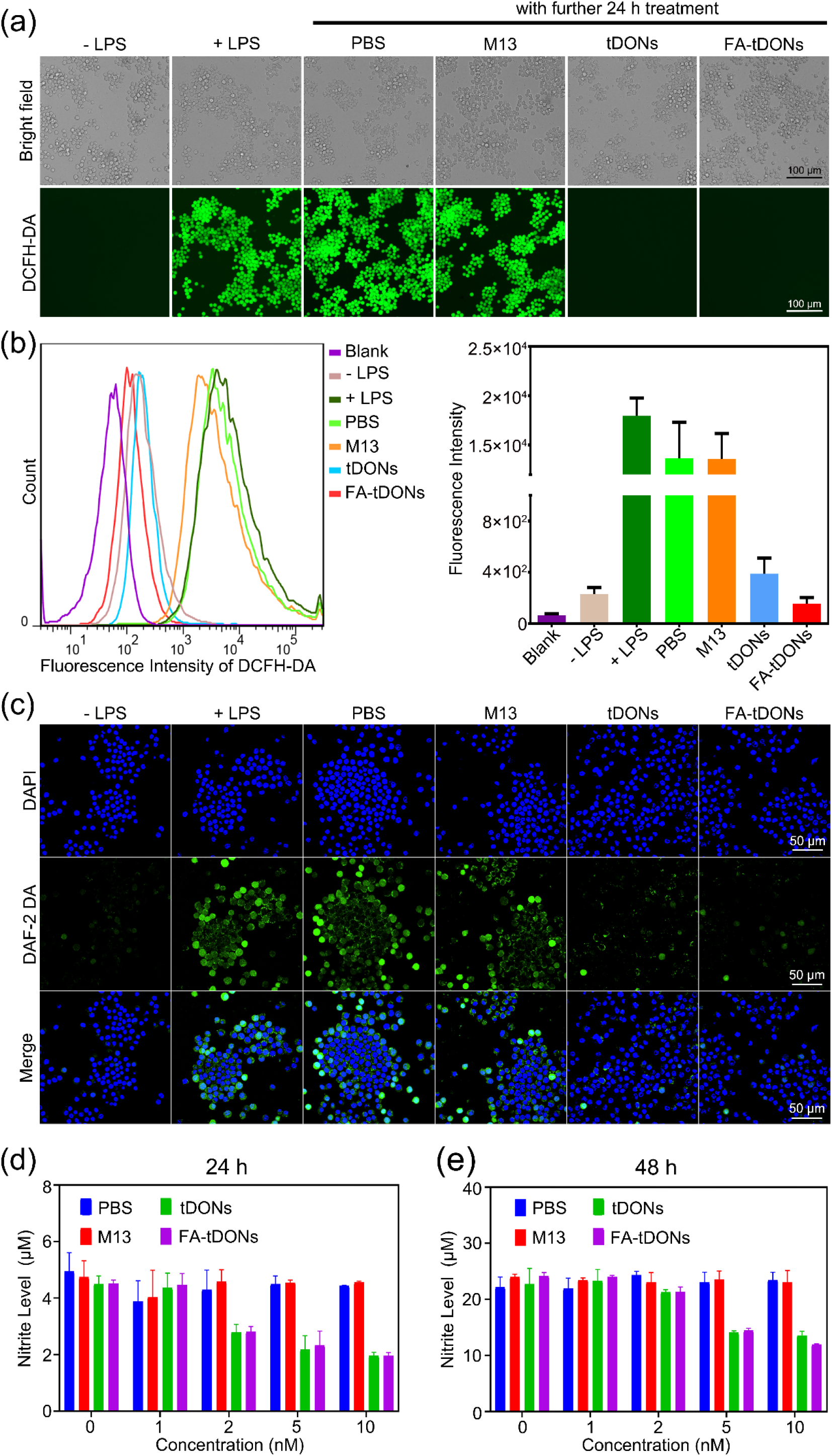
ROS and endogenous NO scavenging ability of FA-tDONs in M1 macrophages. (a) Fluorescence images and (b) flow cytometry were used to detect intracellular ROS levels by the ROS fluorescent probe DCFH-DA. (c) Quantified fluorescence intensity from (b). (d) Confocal microscopy images of RAW 264.7 cells with different treatments. NO and nuclei were stained with DAPI (blue) and DAF-2 DA (green), respectively. Production of nitrite at (e) 24 h and (f) 48 h after treatment with LPS and different concentrations of tDONs and FA-tDONs.

Activated macrophages also secrete high levels of endogenous NO to promote the progression of RA. We first visualized the intracellular NO levels as detected by DAF-2 DA. High fluorescence intensity of NO was observed in RAW 264.7 cells activated with LPS, but the intensity significantly decreased after the addition of tDONs and FA-tDONs for 24 h (Figure 3d), indicating their efficient removal of NO. Next, intracellular NO levels were quantitatively measured by Griess assay. 5 nM tDONs and FA-tDONs could significantly decrease the the NO level after 24 h of treatment (Figure 3e), and showed a long-lasting NO scavenging effect after 48 h of treatment (Figure 3f).

### In vitro M1 to M2 phenotypic transition of macrophages

Strategies for scavenging ROS and NO have been employed to regulate macrophage phenotypes. After confirming the strong ROS and endogenous NO scavenging activity of FA-tDONs in M1 macrophages, we evaluated the in vitro M1 to M2 phenotypic transition by FA-tDONs. Western blot analysis revealed that FA-tDONs decreased the expression of iNOS (an M1 marker), while the expression levels of CD206 (a representative M2 marker) significantly increased (figure 4a). Immunofluorescence staining exhibited a similar tendency (figure 4b), indicating the successful repolarization of macrophages from the M1 to M2 type.

**Figure 4.**
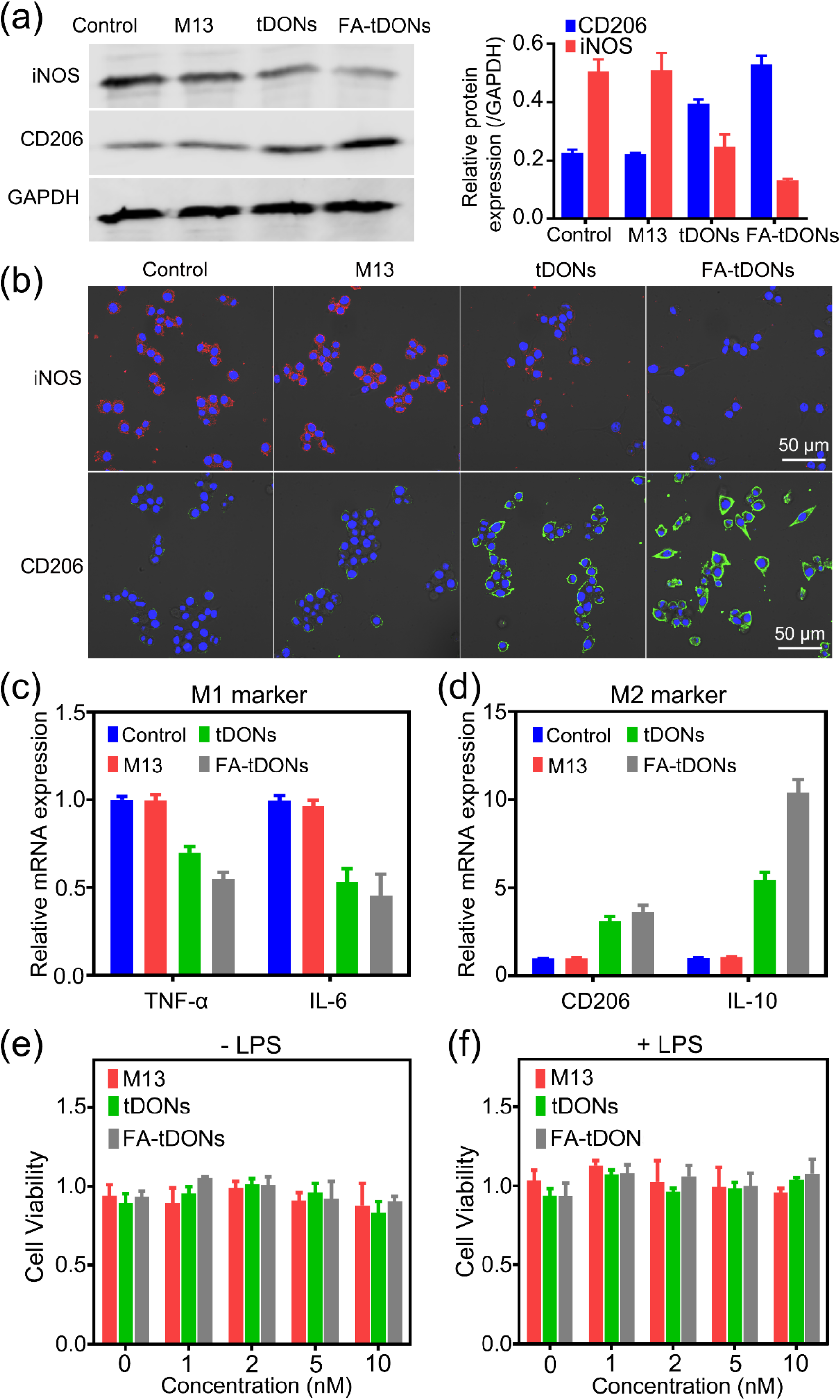
In vitro M1 to M2 phenotypic transition of macrophages induced by tDONs and FA-tDONs. (a) Western blot analysis of the protein expression of iNOS (M1 macrophage marker) and CD206 (M2 macrophage marker). (b) Immunofluorescence staining of iNOS (red), CD206 (green) and nuclei (blue). qRT–PCR analysis of the mRNA expression of (c) M1 macrophage markers (TNF-α and IL-6) and (d) M2 macrophage markers (CD206 and IL-10). Viability of RAW264.7 cells (e) without or (f) with LPS stimulation after incubation with M13, tDONs or FA-tDONs at various concentrations for 48 h (n = 3).

Moreover, the mRNA expression of macrophage markers, including TNF-α, IL-6, CD206 and IL-10 (figure 4c and 4d), was detected by real-time fluorescent quantitative PCR (qPCR). Similarly, FA-tDONs significantly reduced the mRNA expression levels of proinflammatory cytokines TNF-α and IL-6. In contrast, anti-inflammatory M2 markers CD206 and IL-10, were remarkably increased after FA-tDON treatment. Notably, we observed that tDONs also showed a macrophage polarization effect but to a lesser extent, which might be due to their lower cellular internalization efficiency. Taken together, FA-tDONs can modulate macrophage polarization and subsequently exert anti-inflammatory activity by synergistically scavenging ROS and endogenous NO. More importantly, tDONs and FA-tDONs did not exhibit cytotoxicity for 48 h at concentrations up to 10 nM, as determined by CCK 8 assay (figure 4e and 4f). The good biocompatibility of FA-tDONs further motivated us to explore their potential biomedical applications in vivo.

### In vivo targeting ability and therapeutic efficacy in CIA mice

Cy5.5-labeled FA-tDONs were intravenously administered to healthy and collagen-induced arthritis (CIA) model mice to evaluate their targeted accumulation. Arthritis was well developed four weeks after one immunization according to previously published protocols ^26^(Figure 5a) and characterized by obvious paw swelling and redness (Figure 5b). Intravenous injection of FA-tDONs led to a prominent signal in the inflamed joints of CIA mice, and the signal peaked at 1 h and lasted for at least 24 h post-injection (figure 5c and 5d). In contrast, the fluorescence intensity in the noninflamed joints of normal mice was much weaker, which was hardly detectable at 24 h post-injection. In addition, in one CIA model mouse, one paw was morbid, and the other was normal. We also observed preferential accumulation of FA-tDONs in the inflamed joints rather than the normal paw (figure S7), mainly due to active targeting delivery by FA modification and the passive accumulation by ELVIS effect.

**Figure 5.**
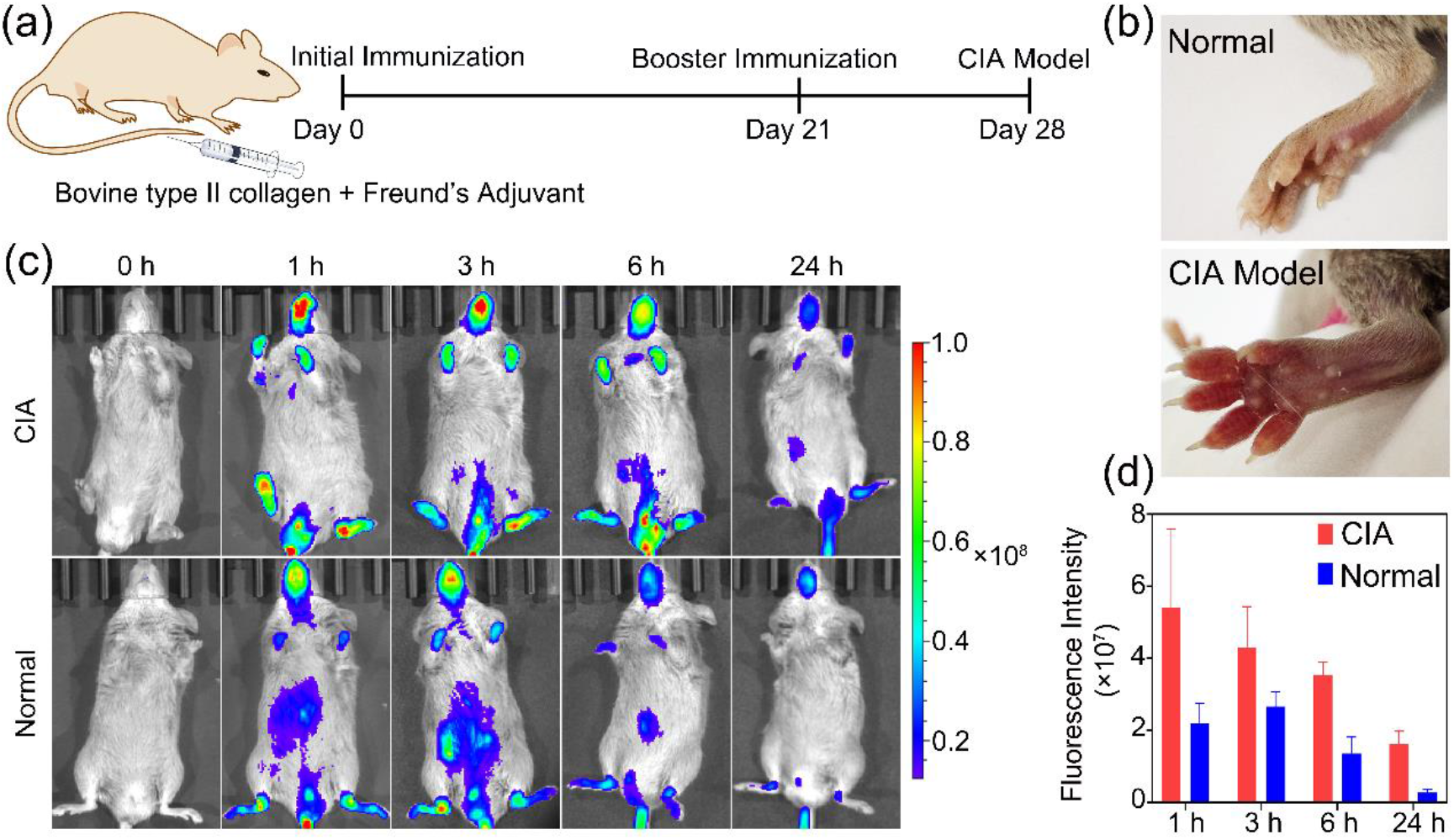
In vivo targeting of FA-tDONs. (a) Modeling procedure of collagen-induced arthritis (CIA) model mice. (b) Twenty-eight days after the first immunization 28 days, photographs of the foot joints of normal and CIA model mice showed evident redness and swelling in CIA model mice. (c) Image of mice after intravenous injection of FA-tDONs into the CIA model and normal mice. (d) Fluorescence intensity quantification from (c).

The CIA mice were then randomly divided into four groups and injected intravenously with saline, m13, tDONs, and FA-tDONs every other day for 22 days (figure 6a). Among all the groups, the mice subjected to FA-tDONs showed little paw swelling as well as much less redness, which was similar to the healthy mice (figure 6b). Bone erosion in all groups was also explored by micro-CT. Obvious ankle erosion and rough bone surfaces were observed in the saline- and M13-treated groups, whereas the FA-tDON-treated group displayed an improvement. Furthermore, histological examination of the paws was carried out to evaluate the biological effects, and the FA-tDON group exhibited the mildest inflammation with smooth articulation of the cartilage surface and little synovial hyperplasia. Next, the arthritis severity in the clinical assessment was determined by the clinical arthritis score. The FA-tDON group achieved significantly decreased clinical scores and better end outcomes than the others (figure 6c and 6d). Specific immunohistochemistry staining and quantitative analysis were then used to measure the expression of the proinflammatory cytokines NF-α and IL-6, and FA-tDON treatment reduced them close to normal levels (figure 6e and 6f). Furthermore, as measured through immunofluorescence staining and quantitative analysis (figure 7g and 7h), the expression of M1 macrophage-specific biomarker iNOS in inflamed joints was remarkably elevated compared with that in healthy mice, but FA-tDON treatment significantly reduced its level, accompanied by the upregulation of anti-inflammatory M2 marker CD206.

**Figure 6.**
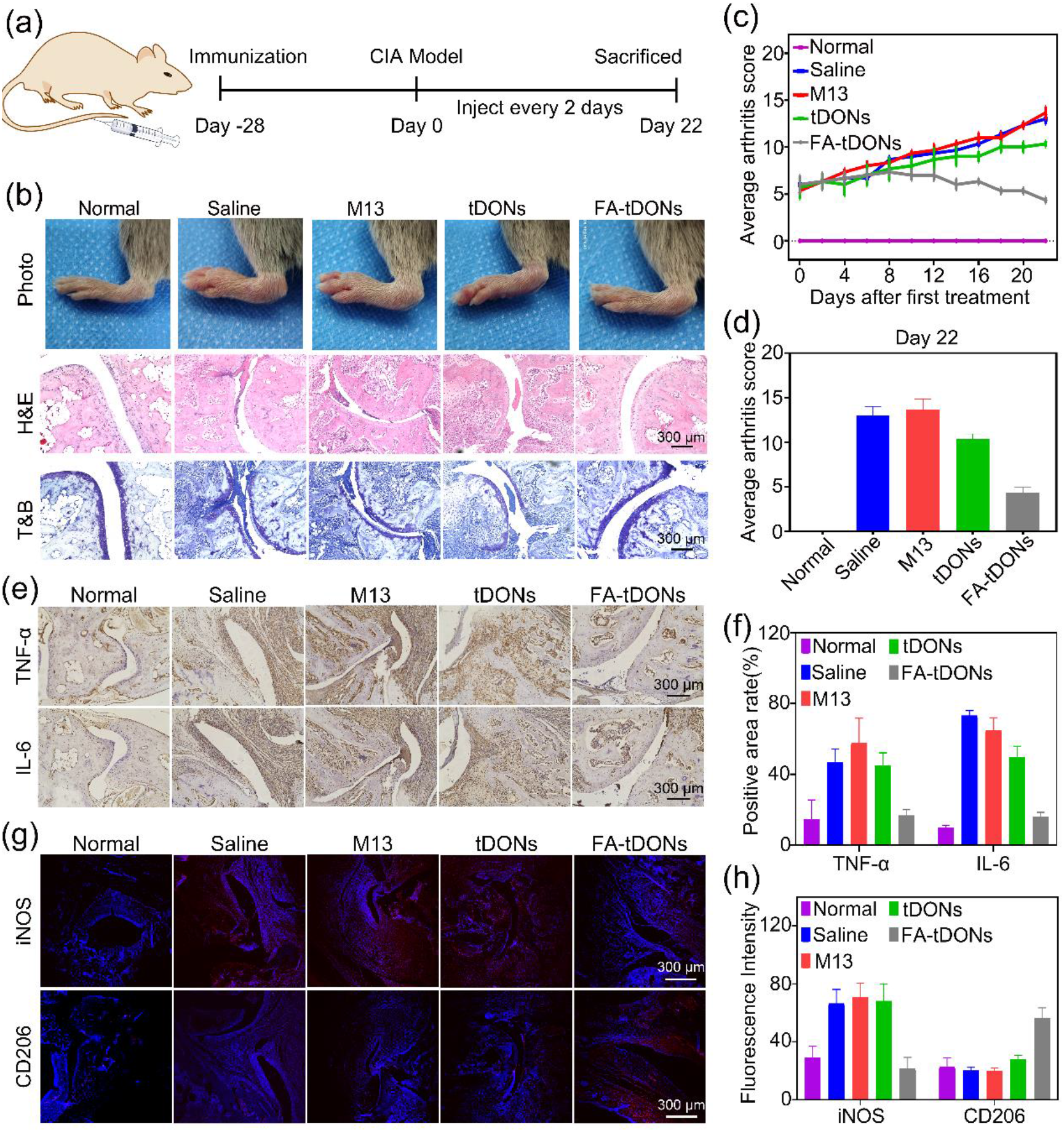
In vivo therapeutic efficacy of FA-tDONs. (a) Overall experimental timeline for the in vivo experiment with intravenous injections every two days for 22 days. (b) Representative photographs, microcomputed tomograph, hematoxylin-eosin (H&E) staining and toluidine blue (T&B) assay of inflamed joints. (c) The every-other-day clinical score and (d) the end-point clinical score of CIA model mice treated with different groups. (e) Immunohistochemical staining of TNF-α and IL-6 in inflamed joints. (f) Quantitative analysis of TNF-α and IL-6 expression levels in inflamed joints from (e). (g) Immunofluorescence staining of iNOS (M1 macrophage marker) and CD206 (M2 macrophage marker) in inflamed joints. (h) Quantitative analysis of iNOS and CD206 expression levels in inflamed joints from (g).

**Figure 7.**
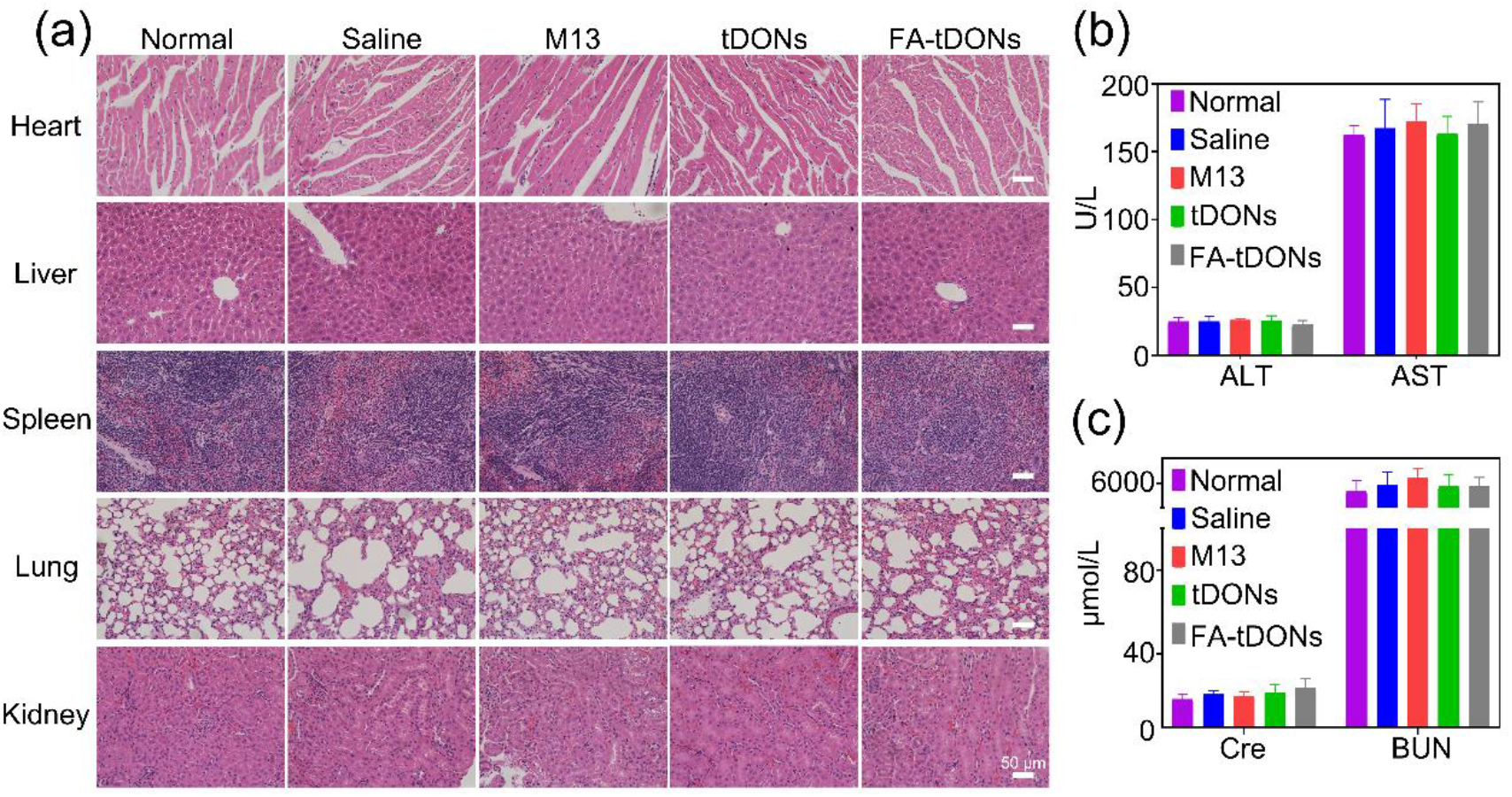
In vivo safety evaluation of FA-tDONs. (a) Images of H&E-stained sections of major organs from CIA model mice at the end of therapy. (b) Blood biochemistry analysis of liver function indexes (ALT and AST) and (c) renal function indexes (Cre and BUN) at the end-point of therapy. Abbreviations are ALT: alanine transaminase; AST: aspartate transaminase; BUN: blood urea nitrogen; CREA: creatinine.

### Safety evaluation of FA-tDONs in vivo

Major organ histology and blood biochemistry analysis of mice after different treatments were performed to evaluate the potential toxicity of FA-tDONs in vivo. H&E staining of the major organs indicated no obvious histological changes in all treatment groups compared to the saline control (figure 7a). Meanwhile, there was no significant difference among groups in liver function indexes alanine transaminase (ALT) and aspartate aminotransferase (AST) and renal function indexes blood urea nitrogen (BUN) and creatinine (Cre) (figure 7b and 7c), suggesting no toxicity of FA-tDONs to the liver and kidney. Collectively, these results implied that FA-tDONs were a highly safe and effective platform for RA therapy.

## Conclusions

In summary, we explored the potential of DNA origami nanostructures as a nanomedicine for RA therapy. In this work, the triangle DNA origami nanostructures exhibit effective ROS and NO scavenging abilities, which are attributed to the polarization of proinflammatory M1 macrophages into anti-inflammatory M2 macrophages. At the mouse level, the triangle DNA origami nanostructures decrease the expression of the proinflammatory cytokines TNF-α and IL-6, and alleviate RA disease progression. One one hand, due to the existence of ELVIS effect, the rationally designed nanosized particles can be passively accumulated at RA sites. On the other hand, this nanomedicine can also actively target to the arthritic sites because of the modification of folic acid, increasing its therapeutic efficacy. Our RA nanomedicine also features precise structures, long duration, high biosafety and biostability. Though DNA origami nanostructures are excellent drug delivery carriers, this work demonstrated for the first time that they possess the potential as nanomedicine themselves for RA treatment.

## Acknowledgements

We thank the National Key Research and Development Program of China (2019YFA0900400), National Natural Science Foundation of China (21977046), Jiangsu Province Key Research and Development Program: Social Development Project (BE2021653), Program for Innovative Talents and Entrepreneur in Jiangsu, and Fundamental Research Funds for the Central Universities for financial support.

